# Evolution-assisted engineering of formate assimilation via the formyl phosphate route in *Escherichia coli*

**DOI:** 10.1101/2025.10.13.681875

**Authors:** Jenny Bakker, Maximilian Boinot, Karin Schann, Jörg Kahnt, Timo Glatter, Tobias J. Erb, Maren Nattermann, Sebastian Wenk

## Abstract

The transition towards a sustainable bioeconomy requires the use of alternative feedstocks, with CO_2_-derived formate emerging as a promising candidate for industrial biotechnology. Despite its beneficial characteristics as a feedstock, microbial assimilation of formate is limited by the inefficiency of naturally evolved formate-fixing pathways. To overcome this limitation, synthetic formate reduction cascades could enable formate assimilation via formaldehyde, a key intermediate of several existing one carbon assimilation pathways. Recently, the formyl phosphate route, combining ATP-dependent activation of formate to formyl phosphate, followed by its reduction to formaldehyde, was developed through enzyme engineering and characterized *in vitro*. In this work, we successfully established the formyl phosphate route *in vivo* by developing a selection strategy that couples formate reduction to growth in a threonine/methionine auxotrophic *Escherichia coli*. Through adaptive laboratory evolution, we achieved formate-dependent growth via this novel pathway. Evolved strains were capable of growing robustly with formate concentrations between 20 mM and 100 mM with glucose in the co-feed. Genomic and proteomic analyses together with activity assays uncovered that formate activation was limiting *in vivo*. This discovery guided the rational engineering of a strain capable of efficient formate assimilation through the formyl phosphate route. By demonstrating that novel enzyme activities can link formate reduction to cell growth, our study shows how synthetic metabolic routes can be functionally established inside the cell, paving the way for the engineering of more complex synthetic pathways.

## Introduction

The one-carbon (C1) molecule formate is a highly promising feedstock for a sustainable CO_2_-based bioeconomy (Yishai et al., 2016). It can be efficiently produced from CO_2_ via photoreduction (Pan and Heagy, 2020), electrochemical reduction (Ewis et al., 2023) or hydrogenation (Saeidi et al., 2021) and is highly soluble in water, avoiding gas-liquid mass transfer limitations prevalent in gas fermentation (Cotton et al., 2020).

Recent progress has enabled the engineering of model microbes to grow on formate (Bang et al., 2020; Kim et al., 2020; Wenk et al., 2022). Additionally, natural formatotrophs have been engineered for biotechnological production with some success (Chang et al., 2022; Yoon et al., 2021). Under aerobic conditions, formate assimilation relies exclusively on the enzyme formate-tetrahydrofolate ligase (FTL), which catalyzes a two-step reaction: a formate kinase reaction followed by the formylation of tetrahydrofolate (THF) to produce formyl-THF. This intermediate is then reduced to methylene-THF (mTHF) and incorporated into biomass (Bar-Even, 2016). Consequently, current formate assimilation strategies are largely restricted to mTHF-dependent pathways, such as variants of the serine cycle and the reductive glycine pathway (Bar-Even, 2016). This limited availability of formatotrophic/formate assimilation pathways constrains the potential of formate as a sustainable substrate for the bio economy (Bar-Even, 2016).

To expand the options for formate assimilation, formaldehyde could be used as an intermediate instead of mTHF. Formaldehyde is primarily assimilated via the ribulose monophosphate (RuMP) cycle and the xylulose monophosphate (XuMP) cycle, which are native to methylotrophic bacteria and yeasts (Chistoserdova et al., 2009; Pham et al., 2022; Wefelmeier et al., 2023; Wegat et al., 2022). In addition to these natural pathways, synthetic formaldehyde assimilation routes have been designed and implemented, including the homoserine cycle (He et al., 2020), the formolase pathway (Siegel et al., 2015), the EuMP cycle (Wu et al., 2023), the SACA pathway (Lu et al., 2019) or the FORCE pathway (Chou et al., 2021). By establishing a metabolic route from formate to formaldehyde and expressing this route in natural or synthetic methylotrophs growing via the above pathways, these organisms could be converted into formatotrophs. This strategy would significantly expand the potential of formate as a renewable carbon substrate, unlocking formate-based production of value-added chemicals.

To catalyze the energetically challenging conversion of formate to formaldehyde, two synthetic pathways have been proposed: formate activation either through phosphorylation or formylation of coenzyme A (CoA), followed by a reduction of the activated intermediate (Bar-Even, 2016; Nattermann et al., 2023) (**Figure 1**). Previous studies have focused primarily on the CoA-dependent route (Hu et al., 2022; Wang et al., 2021). However, this pathway is highly energy-intensive, requiring 2 ATP per formate reduction, and its reliance on CoA as a cofactor poses a risk of perturbing the cellular CoA pool, potentially leading to metabolic interference. In contrast, the formyl phosphate route requires only one ATP and has little overlap with extant metabolism. This second formate reduction route was recently established by combining the promiscuous formate kinase activity of *E. coli* acetate kinase (ACK) with an engineered formyl phosphate reductase (FPR), the latter being a triple mutant of the promiscuous N-acetyl-gamma-glutamyl reductase from *Denitrovibrio acetiphilus* (DaArgC S178V G182V L233I) (Nattermann et al., 2023). Although the functional activities of both pathways have been demonstrated both *in vitro* and *in vivo*, bacterial growth supported by either route has yet to be achieved.

**Figure 1:**
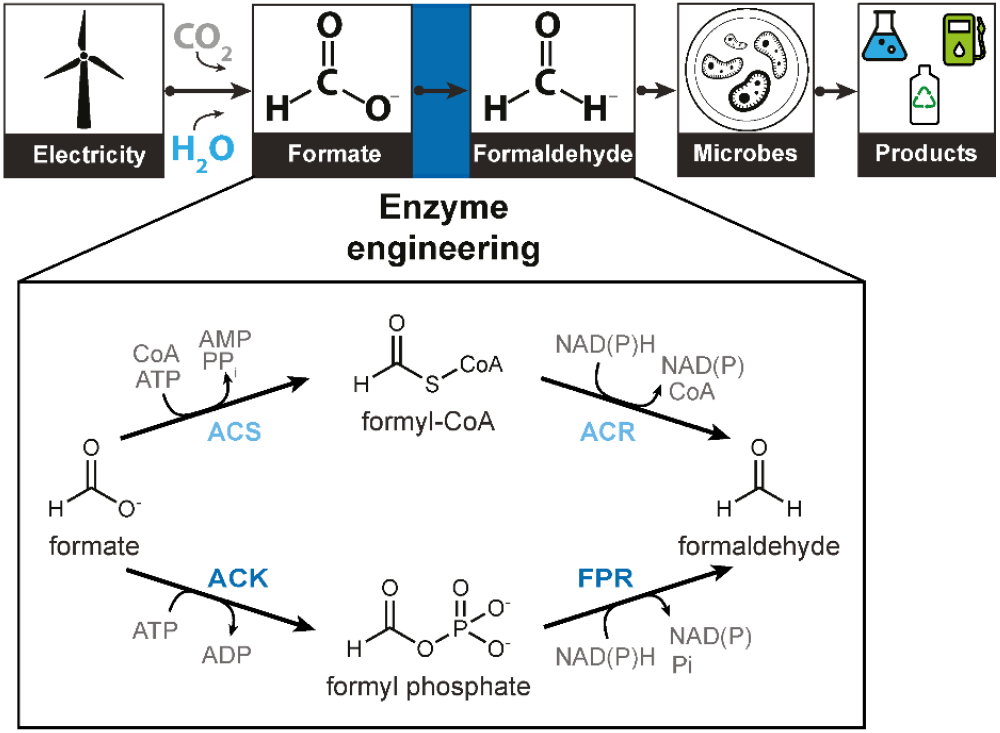
Synthetic formate reduction cascades in a formate bioeconomy. The formate bioeconomy concept envisions the conversion of carbon dioxide into formate using renewable energy. Formate acts as a versatile feedstock for microbial production of value-added chemicals. Using synthetic formate reduction cascades, formate is assimilated into biomass through an intermediate, formaldehyde, and subsequently utilized for bioproduction. The reduction of formate to formaldehyde requires activation via either formyl-CoA or formyl phosphate intermediates, which are then reduced to formaldehyde. Key enzymes involved in these pathways include ACS – acetyl-CoA synthetase, ACR – acyl-CoA reductase, ACK – acetate kinase and FPR – formyl phosphate reductase.

In this study, we successfully implemented the formyl phosphate route in *E. coli*, demonstrating that the synthetic cascade can be used for metabolic incorporation of formate. Utilizing a specifically engineered biosensor strain that requires formaldehyde for the biosynthesis of threonine and methionine, we established the pathway through a combination of engineering and adaptive laboratory evolution (ALE). Genomic and proteomic analyses of the evolved strains revealed that during evolution, plasmidic mutations had led to increased ACK expression thereby optimizing the *in vivo* activity of the formate reduction cascade. These findings allowed us to precisely engineer a strain with ACK and FPR levels tailored to efficient, growth-coupled formate reduction. This work demonstrates that synthetic formate reduction cascades can sustain flux sufficient for metabolic complementation, a prerequisite for full synthetic formatotrophy.

## Results

### Adaptive laboratory evolution enables *E. coli* growth via the formyl phosphate route

To test whether the formyl phosphate route could support growth of *E. coli*, we devised a strategy to couple formate reduction to cell growth. Previously, we designed and constructed growth-coupled formaldehyde biosensor strains that depend on formaldehyde assimilation for cell growth and can detect a wide range of intracellular formaldehyde concentrations (Schann et al., 2024). Out of the different sensor strains available, we chose the 4-hydroxy 2-oxobutanote (HOB) biosensor strain for implementation of the formyl phosphate route. The HOB biosensor strain is highly sensitive to formaldehyde and can grow at concentrations as low as 32 µM, which aligns with the productivity of the cascade (up to 50 µM/h *in vivo*) (Nattermann et al., 2023). At the same time, it can tolerate formaldehyde concentrations that are toxic to wild-type *E. coli* (up to 2 mM) thanks to an engineered formaldehyde assimilation module, which serves as a sink for formaldehyde, thus mitigating its toxicity (**Supplementary Figure1**). Selection in the HOB biosensor strain is based on an engineered threonine/methionine auxotrophy created by the deletion of aspartate-semialdehyde dehydrogenase (Δ*asd*), which is relieved through the assimilation of formaldehyde and pyruvate into homoserine (and subsequently threonine and methionine) via the HAL/HAT formaldehyde assimilation module of the synthetic homoserine cycle (He et al., 2020) (**Figure 2A**). As threonine and methionine account for ∼5% of the cellular biomass (Simensen et al., 2022), the strain relies on formate reduction to formaldehyde to generate this fraction of biomass, while the remaining biomass is derived from glucose.

**Figure 2:**
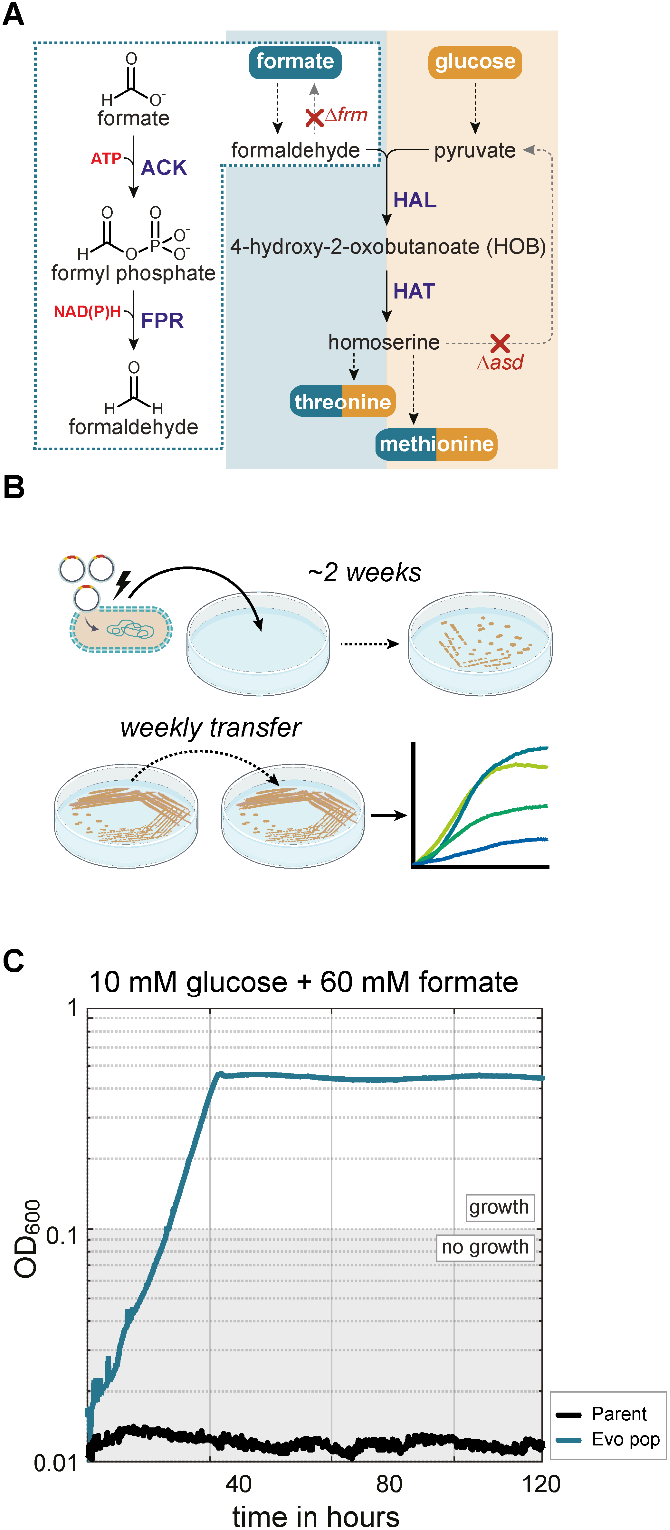
Formate reduction is enabled through direct selection and plate evolution. (A) Metabolic scheme of the HOB biosensor strain, which was used to test the *in vivo* activity of the formyl phosphate route. In this strain, formate is converted to formaldehyde via the combined activities of ACK and FPR. The formaldehyde is further condensed with pyruvate into homoserine through the activities of HAL and HAT. Homoserine is subsequently converted into threonine and methionine, relieving the auxotrophies of the strain. (B) Schematic overview of the adaptive laboratory evolution experiment conducted to achieve formate-dependent growth. Plasmids encoding the ACK-FPR operon were introduced into the HOB biosensor strain via electroporation and plated on minimal medium plates containing 10 mM glucose and 60 mM formate (selective medium). Small colonies appeared after two weeks and were re-streaked weekly on fresh plates. (C) Formate-dependent growth of the evolved population. The evolved population, which consistently grew on selective plates, was cultivated in selective liquid medium. Growth characterization demonstrated formate-dependent growth, as shown by the blue line. Abbreviations: ACK – acetate kinase; FPR – formyl phosphate reductase; HAL – HOB aldolase (catalyzed by *E. coli* 2-keto-3-deoxy-L-rhamnonate aldolase RhmA); HAT – HOB aminotransferase.

Before testing the formyl phosphate route *in vivo*, we modified the HOB biosensor strain to remove enzymes involved in acetate production from acetyl-CoA (Δ*ackA-pta*), to both generate a clean background for ACK production and avoid draining of formyl phosphate into formyl-CoA by PTA. For the expression of the formyl phosphate route, we chose a plasmid-based strategy, as it offers greater flexibility for transformation, tuning of expression levels, exchange of regulatory elements, and adaptive evolution. We cloned the candidate genes for ACK (*E. coli ackA*) and FPR (*Denitrovibrio acetiphilus argC S178V G182V L233I*) into a synthetic operon. In the operon, expression of ACK was controlled by a ribosomal binding site (RBS) of medium strength (rbsC) and expression of FPR by a strong RBS (rbsA) (Zelcbuch et al., 2013), as FPR was determined to be rate-limiting for the cascade *in vitro* (Nattermann et al., 2023). We transformed the HOB biosensor strain with the plasmid recovering the cells on non-selective medium. Thereafter, we tested the growth of transformed strains in selective medium with formate and glucose as carbon sources. However, the cells were unable to grow.

As no growth was achieved via the standard transformation procedure, we decided to directly selected for *in vivo* pathway activity after transformation (**Figure 2B**). For this, the HOB biosensor strain was electroporated with the plasmids encoding the formate reduction pathway, followed by plating on minimal medium plates supplemented with glucose and formate. The appearance of colonies on these plates after several days of incubation signaled a functional formate reduction cascade, allowing further cultivation to enhance *in vivo* activity. To enhance growth, these colonies were transferred to fresh plates weekly over approximately two months (**Supplementary Figure 2**). We then tested the growth characteristics of the adapted populations in liquid culture. As shown in **Figure 2C**, the evolved population demonstrated efficient growth in formate-containing medium, while the original strain did not grow. This result indicated that the strains developed a pathway-related adaptation enabling formate-dependent growth.

### Plasmid evolution adapts protein levels to enable formate reduction

After observing formate-dependent growth in the evolved population, we isolated single colonies and conducted growth experiments with the evolved strains (Evo1 and Evo2). Both strains exhibited formate-dependent growth but displayed distinct growth phenotypes with Evo1 growing faster and with lower formate concentrations compared to Evo2 (**Figure 3A and Supplementary Figure 3**). This can be attributed to better formaldehyde production by Evo1 which we observed during growth using the NASH assay (**Supplementary Figure 3)**.

**Figure 3:**
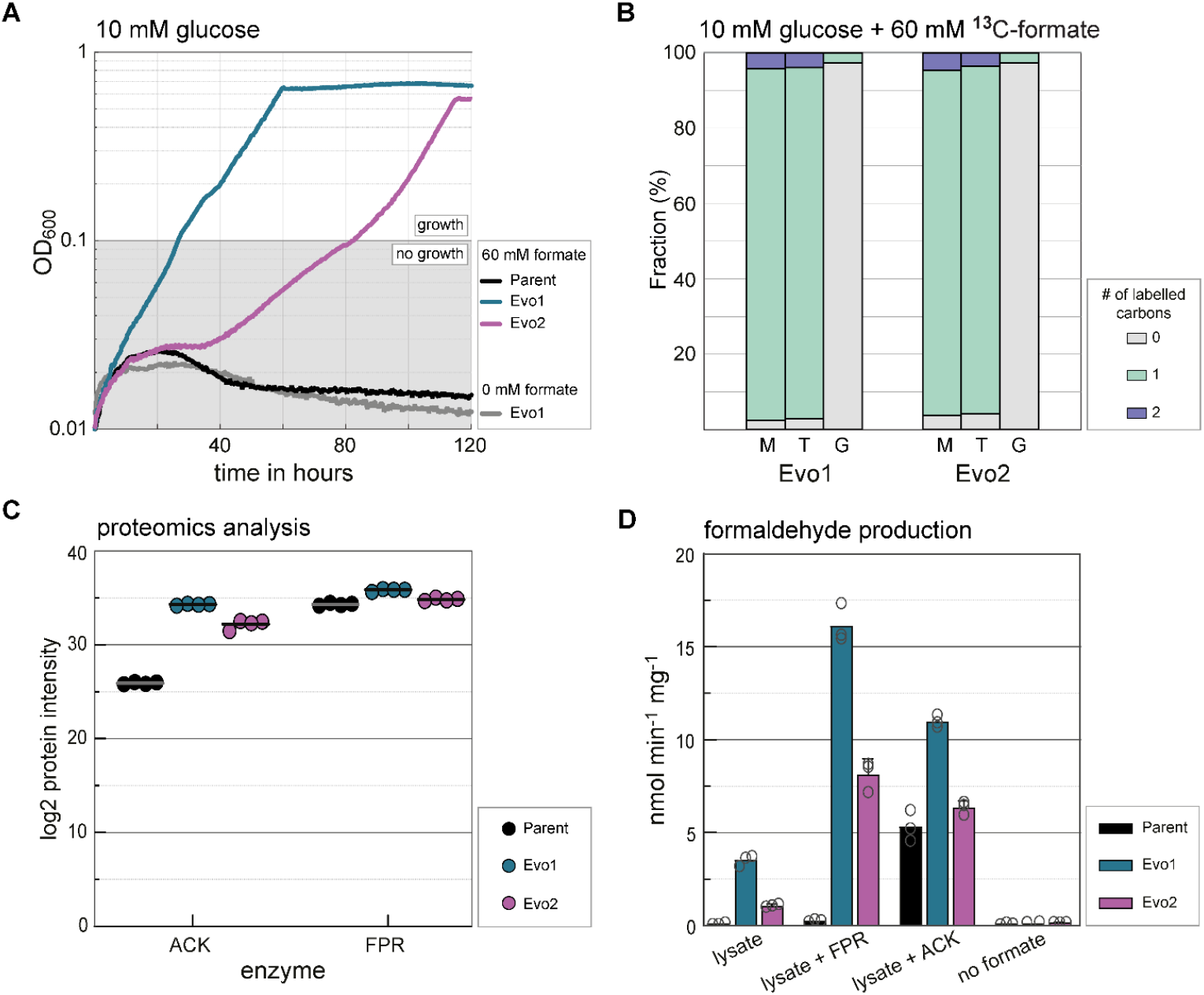
Adaptation of protein levels enables formate assimilation via the formyl phosphate route. (A) Growth characterization of two isolates from the evolved population. Isolates Evo1 and Evo2 were cultivated in selective medium, and their growth phenotypes were compared to the unevolved (parent) strain. Evo1 exhibited faster growth and reached a higher maximum OD_600_ than Evo2. (B) ^13^C-labelling confirms formate reduction to formaldehyde *in vivo*. Incorporation of ^13^C-labelled formate into threonine and methionine was observed in both isolates (turquoise bars), validating the activity of the formyl phosphate route. (C) Proteomic analysis revealed protein-level adaptations during evolution. ACK levels were highly elevated in both isolates compared to the parent strain, with Evo1 displaying higher levels than Evo2. Minor changes in FPR levels were observed between the parent and evolved isolates. (D) Enzymatic activity assays of ACK and FPR. Formaldehyde production was elevated in the lysates of Evo1 and Evo2 (turquoise and purple bars) compared to the parent strain (black bar). The addition of purified ACK or FPR to the lysates revealed that the parent lysate was deficient in ACK. T – threonine; M – methionine and G – glycine.

To confirm the activity of the formyl phosphate route, we conducted labelling experiments with ^13^C formate. Based on the pathway reactions and the metabolic architecture of the HOB biosensor strain, the ^13^C label of formate should end up first in ^13^C-formaldehyde, which is subsequently condensed with pyruvate to form single-labeled ^13^C-homoserine. Finally single labeled ^13^C-threonine and ^13^C-methionine should be produced from ^13^C-homoserine. Following growth of the evolved strains with ^13^C-formate, we extracted their proteinogenic amino acids and analyzed them using liquid chromatography/mass spectrometry (LC-MS). The labeling pattern matched the predictions, confirming the *in vivo* activity of the formyl phosphate route (**Figure 3B**).

To explore the evolutionary mechanisms enabling growth of the evolved strains, we investigated whether formate-dependent growth was dependent on the pathway-encoding plasmid. To achieve this, we conducted two experiments: first, curing Evo1 of its plasmids, and second, introducing the plasmid isolated from Evo1 into a naïve HOB biosensor strain. Growth experiments with these strains revealed that strains carrying the plasmid from Evo1 (both Evo1 and the naïve HOB biosensor strain transformed with the Evo1 plasmids) exhibited growth whereas the plasmid-cured Evo1 did not, indicating that plasmid-encoded genes are essential for formate reduction *in vivo* (**Supplementary Figure 4**). To determine whether the plasmids themselves had evolved, we performed Nanopore sequencing to analyze the complete plasmid structure (Kasianowicz et al., 1996; Zheng et al., 2023). The sequencing confirmed plasmid evolution, revealing plasmid multimerization (**Supplementary Figure 4**) and mutations in ACK. Evo1 showed a point mutation in the RBS, while Evo2 displayed a 4 bp insertion in *ackA*, altering the start codon and removing its polyhistidine-tag. Whole genome sequencing of the evolved strains did not show any mutations compared to the unevolved parent strain (**Supplementary Table 2**) and as growth of the naïve HOB biosensor strain was achieved upon transformation with the evolved plasmids, we focused our subsequent analysis on plasmid-derived evolutionary effects.

The observed plasmid mutations led us to investigate their influence on the expression of pathway enzymes. To this end, we conducted a comparative proteomic analysis on parent and evolved strains. The results of this experiment showed that the evolved strains expressed dramatically higher levels of ACK compared to the parent strain (increased by 3 orders of magnitude), whereas FPR levels, already high in the parent strain, were less affected by evolution (**Figure 3C**). This finding indicates that, in the original configuration, ACK expression was too low to generate the metabolic flux from formate to formaldehyde required for cell growth.

To investigate this hypothesis, we tested formaldehyde production from formate in cell lysates of all strains. The results confirmed our hypothesis: detectable amounts of formaldehyde were produced only in the evolved strains (**Figure 3D**). We also observed a correlation between growth and formaldehyde production, with the superior growth of Evo1 mirrored by elevated formaldehyde production in its lysate (**Figure 3A and 3D**).

To test whether the low ACK level in the parent strain is responsible for its lack of formaldehyde production (and thus growth), we supplemented the cell lysates with either purified ACK or FPR. In the parent strain lysate, ACK supplementation significantly increased formaldehyde production, while FPR supplementation had almost no effect on it (**Figure 3D**). In the lysates of the evolved strains, the addition of either enzyme improved formaldehyde production, with FPR addition showing the biggest effect especially for Evo1 which has the highest ACK expression of all three strains. These results confirm that low ACK levels were the primary reason for the lack of growth of the parent strain.

While FPR is the kinetically inferior enzyme within the cascade, with a turnover number two orders of magnitude lower than that of ACK (Nattermann et al., 2023) and a high Km for formyl phosphate (∼20 mM), it seems that high ACK activity is needed to produce sufficient amounts of formyl phosphate. We therefore hypothesize that FPR is undersaturated in the parent strain, making its turnover highly responsive to the concentration of ACK (**Supplementary Figure 5**). This puts evolutionary pressure on the production of both enzymes, explaining why ACK production increased in the evolved strains.

Finally, as the product of ACK, formyl phosphate is instable in aqueous solutions (0.3 – 1%/min in aqueous solution (Nattermann et al., 2023)), its hydrolysis could negatively impact pathway performance. To test for this, we performed an *in vitro* experiment under substrate limitation. As this experiment showed stoichiometric production of formaldehyde from ATP, hydrolysis does not seem to impact pathway performance (**Supplementary Figure 6**).

### Evolution-assisted design of a functional formyl phosphate route

Equipped with a better understanding of the cascade based on our proteomics and *in vitro* analysis, we designed strains with more balanced enzyme levels. Primarily, this meant increasing ACK production, which had been the sole block to growth in the parent strain.

We changed its RBS from medium to strong (rbsA) and integrated the gene into a highly active region of the *E. coli* chromosome (Bassalo et al., 2016). To modulate FPR levels, we expressed it from either the same plasmid used in the initial design (p15A, ∼10 copies) or a higher-copy plasmid (pMB1, ∼20 copies), resulting in two rationally engineered strains: Formate Reducer 1 (FR1) and Formate Reducer 2 (FR2) (**Figure 4A**). In growth experiments, both strains exhibited formate-dependent growth, though with distinct growth phenotypes (**Figure 4B**). FR1 grew slower and to a lower OD_600_ than FR2, matching the expected phenotype we would have expected based on the above pathway analysis.

**Figure 4:**
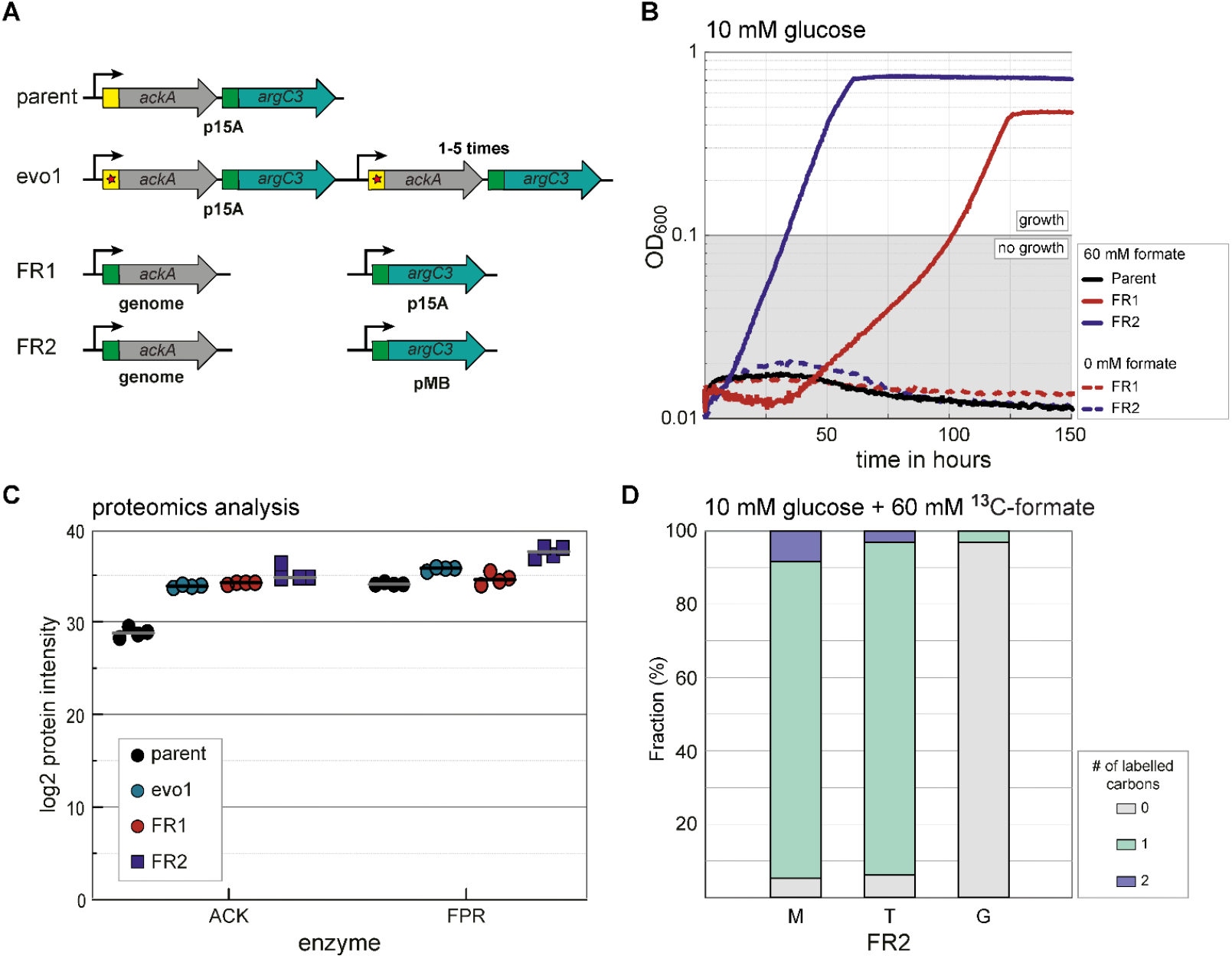
Evolution-inspired engineering enables improved formate reduction *in vivo*. (A) Genetic configurations of ACK (*ackA*) and FPR (*argC3*) in the parent, evolved, and engineered strains. The parent strain expresses the unevolved operon from a plasmid with p15A ori, the evolved strains show multimerization of the same plasmid as well as mutations in the regulatory region of ACK that lead to its upregulation. Reverse engineered strains FR1 and FR2 express ACK with a strong RBS from the chromosome and FPR from either a plasmid with p15A or pMB ori. (B) Growth characterization of engineered strains. The HOB biosensor strain, expressing ACK from the chromosome and FPR from plasmids with either a p15A origin of replication (FR1, red line) or a pMB1 origin of replication (FR2, blue line), was grown in selective medium. FR2 exhibited faster growth and reached a higher final OD_600_ compared to FR1 (C) Proteomic analysis of ACK and FPR levels across the parent, evolved, and engineered strains. ACK protein levels in the engineered strains (FR1, red; FR2, blue) were comparably high as in the evolved strain (Evo1, turquoise). FPR protein levels were consistent across all strains, including the parent strain (black), but were slightly elevated in FR2. (D) Confirmation of formate reduction through ^13^C-labelling. Incorporation of ^13^C-labelled formate into methionine and threonine demonstrated successful formate reduction in the engineered strain. Abbreviations as in figure 3.

Proteomic analysis of the engineered strains showed that ACK levels in FR1 and FR2 closely matched those of Evo1, confirming that the change of RBS combined with the genomic integration achieved the desired effect. As anticipated, FPR levels differed between the strains, with FR1 exhibiting slightly lower FPR levels than Evo1, while FR2 exhibited slightly higher levels than Evo1 (**Figure 4C**). This difference explains the variation in growth phenotypes: FR1 has sufficient ACK for growth, but lower FPR production, whereas FR2 produces more FPR and is thus capable of faster growth. Notably, FR2 produces ACK and FPR at similar levels as Evo1, resulting in comparable phenotypes. Finally, we confirmed the activity of the formyl phosphate route in FR2 through a ^13^C-labeling experiment which showed the same labelling pattern as the evolved strains, confirming the *in vivo* activity of the formyl phosphate route in the engineered strain (**Figure 4D**).

These results highlight that achieving an optimal balance between ACK and FPR expression is crucial for enabling efficient growth via the formyl phosphate route. Notably, our ability to rationally re-engineer this adaptation, guided by evolutionary insights, demonstrates the power of ALE as a pre-screening tool for identifying key genetic modifications that are hard to predict *ab initio*.

## Discussion

In this study, we established formate-dependent growth of the HOB formaldehyde biosensor strain through a synthetic formate reduction cascade. The formyl phosphate route operates via a two-step catalytic process: activation of formate to formyl phosphate via the promiscuous activity of ACK, followed by reduction to formaldehyde by an engineered FPR. Compared to other synthetic formate reduction pathways, the formyl phosphate route is shorter and less ATP-demanding (Bar-Even, 2016; Hu et al., 2022, 2021; Schann and Wenk, 2024).

To establish this route in *E. coli*, we engineered a strain dependent on formaldehyde for cell growth, providing the metabolic framework for growth-coupled selection of the formyl phosphate route (Orsi et al., 2021; Wenk et al., 2018). The HOB biosensor strain depends on formaldehyde assimilation for the biosynthesis of threonine and methionine, representing ∼5% of cell biomass (Simensen et al., 2022). Compared to previous attempts to establish *in vivo* formate reduction (Hu et al., 2022; Wang et al., 2021), our approach uses auxotrophies to make the strain dependent on formate reduction. This design allows for fine-tuned control over the selection pressure, ensuring a more targeted dependency on the synthetic pathway. Furthermore, by providing a metabolic sink for formaldehyde through its assimilation into essential metabolites, this strategy allows the engineered strain to tolerate formaldehyde concentrations that would otherwise be toxic to *E. coli*, thereby mimicking the metabolic architecture of natural methylotrophs.Despite the relatively low selection pressure, our initial strain, which expressed the formate reduction cascade from a plasmid, was unable to grow with formate. To overcome this, we applied ALE on selective plates to facilitate growth via the formyl phosphate route. The choice of solid medium was motivated by the assumption that liquid medium would favor the persistence of cheater cells (non-producers) that exploit freely diffusing formaldehyde produced by evolved cells. A similar observation was reported by Woolston *et al*. when attempting to evolve an optimized methanol dehydrogenase (Roth et al., 2019; Woolston et al., 2018).

Because of its untargeted nature, ALE enables system-wide adaptations that would otherwise be difficult to predict (Dragosits and Mattanovich, 2013). However, the accumulation of multiple mutations across the chromosome can complicate the distinction between beneficial and hitchhiker mutations (Dragosits and Mattanovich, 2013; Phaneuf et al., 2020). Despite these challenges, we were able to identified the key adaptations responsible for activating the formyl phosphate route in the evolved strains.

Two factors likely contributed to this success: (I) Selective plate evolution, which provided more stable cultivation conditions than liquid medium, reducing the accumulation of hitchhiker mutations; and (II) plasmid-based pathway expression, which allowed for the separate analysis of chromosomal and plasmid-borne DNA. This enabled direct plasmid transfer from evolved strains into naïve strains, facilitating the identification of relevant mutations.

Using nanopore sequencing (Kasianowicz et al., 1996; Zheng et al., 2023) and proteomics (Bekker-Jensen et al., 2020), we found that the strains adapted to formate-dependent growth by multimerizing the pathway encoding plasmids and introducing mutations in the regulatory sequences of ACK. These mutations increased ACK expression, achieving an optimal balance between ACK and FPR, thereby enabling efficient conversion of formate to formaldehyde.

These findings challenge our previous understanding of the formyl phosphate route. In earlier work, we strongly expressed both ACK and FPR, observing that the pathway was primarily limited by FPR when tested in cell lysate (Nattermann et al., 2023). This led us to hypothesize that moderate expression of ACK would improve growth-coupled selection by reducing protein burden. However, our current results contradict this assumption, revealing that we had previously underestimated the impact of *K*_m_ on pathway performance. Our new findings demonstrate that elevated levels of ACK are essential to maintain sufficient formyl phosphate production in vivo, allowing FPR to operate at rates compatible with growth. This highlights how limitations in a downstream enzyme can impose stringent upstream requirements, ensuring a steady substrate supply and efficient flux through the cascade.

Building on this refined understanding, we engineered two strains, FR1 and FR2, capable of growth via the formyl phosphate route. These strains, which express ACK from the chromosome and FPR from a plasmid, could function as “FPR biosensors” providing a modular “plug-and-play” platform for enzyme design, engineering or directed evolution campaigns aiming at further improving FPR – a significant improvement over our previous approach in which the screening capacity was limited to the readout of the NASH assay (Nattermann et al., 2023).

This study lays the foundation for enabling full growth of natural or engineered methylotrophs through formate reduction by demonstrating formate-dependent growth via the formyl phosphate route. Further optimization – such as enhancing enzyme expression, improving catalytic efficiency or introducing the enzymes into a synthetic bacterial microcompartment – will be instrumental to fully realize this potential. Here, formaldehyde biosensor strains with higher formaldehyde dependency such as the LtaE or the RuMP biosensor strains previously designed by us could be of great use.

## Acknowledgements

This work was supported by the German Ministry of Education and Research Grant 031B0850B (MetAFor) and the Max Planck Society. The authors are grateful to Arren Bar-Even who initiated the project and was an inspiring mentor to all of us. The authors thank Abdelrahman Alnahhal for his support in the experiments, Saleh Alseekh for performing LC-MS analysis, Kyle Moynahan for helping with data analysis and Ari Satanowski for critical reading of the manuscript and helpful suggestions.

## Author contributions

S.W. conceptualized the project and designed and supervised the research.

S.W., J.B., M.B., K.S. and M.N. wrote the manuscript with contributions from the other authors

S.W., J.B., M.B., and K.S. designed and performed the *in vivo* experiments and analyzed the data.

M.N. performed the *in vitro experiments* and analyzed the data.

T. G. and J. K. performed the LC-MS measurements for the proteomics experiments and analyzed the resulting data.

T.J.E. supervised the *in vitro* work.

## ORCID

## Methods

### Chemicals and reagents

Primers were synthesized at Integrated DNA Technologies (IDT). DNA fragments were amplified with the high-fidelity polymerase PrimeSTAR Max DNA Polymerase from TaKaRa, and colony PCR reactions were performed using DreamTaq polymerase (Thermo Fisher Scientific). FastDigest restriction enzymes and T4 DNA ligase from Thermo Fisher Scientific were used for restriction-based cloning. The kits used for plasmid isolation and PCR cleanup were purchased from Thermo Fischer Scientific. All chemicals used in the plate reader experiments were ordered from Sigma-Aldrich.

### Growth media

For cloning and strain engineering, cultures were grown in lysogeny broth (LB) medium (10 g/L NaCl, 10 g/L Tryptone, 5 g/L Yeast Extract) supplemented with the relevant antibiotic (50 μg/mL kanamycin, 30 μg/mL chloramphenicol or 100 μg/mL streptomycin). Diaminopimelate (DAP, 0.25 mM) was added to the growth media of the HOB biosensor strain (required due to deletion of *asd*). During growth experiments, cells were cultivated in M9 minimal medium (50 mM Na_2_HPO_4_, 20 mM KH_2_PO_4_, 1 mM NaCl, 20 mM NH_4_Cl, 2 mM MgSO_4_ and 100 μM CaCl_2_), supplemented with trace elements (134 μM EDTA, 31 μM FeCl_3_, 6.2 μM ZnCl_2_, 0.76 μM CuCl_2_, 0.42 μM CoCl_2_, 1.62 μM H_3_BO_3_, 0.081 μM MnCl_2_) and the relevant carbon sources. Precultures were grown in relaxing medium, which is M9 minimal medium supplemented with trace elements, 10 mM glucose, 1 mM isoleucine, 1 mM methionine, 1 mM threonine, 0.25 mM DAP, 50 µM MnCl_2_, and the appropriate antibiotic. If not differently stated in the text, the selection medium consisted of M9 minimal medium supplemented with trace elements, 10 mM glucose, 1 mM isoleucine, 60 mM Na-formate, 0.25 mM DAP, 50 µM MnCl_2_, and the appropriate antibiotic.

### Bacterial strains

Plasmid construction was performed using *E. coli* DH5α. The HOB biosensor strain is based on the *E. coli* SIJ488 strain, which is derived from K-12 MG1655 (Jensen et al., 2015). The KEIO collection was used for the preparation of P1 donor lysates for phage transduction (Baba et al., 2006). All strains and plasmids used in this study are listed in **Table S1**.

### Strain engineering

The HOB biosensor strain was constructed as described in (Schann et al., 2024) by modifying *E coli* SIJ 488 using P1 phage transduction and λ-Red recombineering. We additionally deleted the *ackA-pta* operon to avoid the production of acetate, which is a competing substrate of the formate reducing enzymes tested in this study.

To create the strains FR1 and FR2, *ackA* was integrated into the genome of the HOB biosensor strain and overexpressed from SS9 under the control of the P_pgi_-20 promoter and rbsC, using λ-Red recombineering. Antibiotic cassettes were removed by flippase induction, and all integrations were confirmed by colony PCR.

### Synthetic-Operon construction

All genes were codon optimized for *E. coli* K-12, and an N-terminal 6xHis-tag was attached. The genes were transferred to a cloning vector, attaching a synthetic ribosomal binding site upstream of the gene (Zelcbuch et al., 2013). These cloning vectors were used to assemble synthetic operons via restriction and ligation as described in (Wenk et al., 2018). The assembled construct was then transferred to an expression plasmid containing either the p15A or the pMB1 ori, a streptomycin resistance gene and attaches the P_pgi_-20 promoter to the gene or operon of interest. This assembly was performed using the restriction enzymes EcoRI and PstI (Wenk et al., 2018).

### Adaptive laboratory evolution on M9 agar plates

Electroporation at 1800 mV for ∼ 5 ms was used to transform the HOB biosensor strain with a plasmid containing the FOK-FPR operon. Following a one-hour recovery in M9 relaxing medium without antibiotic, the cells were washed three times and resuspended in M9 minimal medium and plated directly on M9 agarose plates supplemented with trace elements, 10 mM glucose, 1 mM isoleucine, 0.25 mM DAP, 50 µM MnCl2, and 60 mM formate to allow for ALE. Growing strains were streaked to new selective M9 plates with and without 60 mM formate weekly for eight weeks.

### Electroporation of plasmids from evolved strains

Plasmids from Evo1 were isolated with the GeneJET Plasmid-Miniprep-Kit from Thermo Fisher Scientific. Isolated plasmids were transformed into the HOB biosensor strain by electroporation at 1800 mV for 5 ms. Strains were recovered in liquid LB containing 0.25 mM DAP for one hour before being plated on LB agar plates with 0.25 mM DAP and 100 mM streptomycin to select for strains carrying the evolved plasmids.

### Plasmid curing from evolved strains

To cure evolved strains of their plasmids, single colonies were inoculated in liquid LB containing 0.25 mM DAP and no antibiotic and were passaged to fresh medium daily. Every five days, single colonies were isolated from each liquid culture, and they were streaked out on plates containing solid LB agar with 0.25 mM DAP and LB with 0.25 mM DAP and 100 mM streptomycin, respectively, to assess whether the strains had lost their plasmids that provided the antibiotic resistance. After detecting colonies that had lost the resistance, we performed a colony-PCR on single colonies growing on plates without antibiotic using primers that target the cargo of the plasmids used in this study and analyzed the results by performing agarose gel electrophoresis on the PCR products.

### Growth experiments

Prior to all growth experiments, the strains were streaked out from the glycerol stock on LB plates (supplemented with the respective antibiotic and 0.25 mM DAP in case of the HOB biosensor strain) and grown overnight at 37 °C. From these plates, precultures in M9 relaxing medium were inoculated for overnight growth at 37 °C and 220 rpm. The cells were harvested (11000 g, 1 min), washed three times with M9 minimal medium, and diluted to a final OD_600_ of 0.01 in the desired medium. In a Nunc 96-well microplates (Thermo Fisher Scientific), 150 µL of the culture was added to each well and the wells were covered with 50 µL of mineral oil (Sigma-Aldrich) to prevent evaporation. The growth experiments were performed in a BioTek Epoch2 plate reader (BioTek Instrument, USA) which measured the optical density after each kinetic cycle. A cycle consists of 12 shaking steps in which 60 seconds linear and 60 seconds orbital shaking were alternated (1 mm amplitude). The OD_600_ was corrected to present the OD_600_cuvette_ by dividing the OD_600_plate_ by the correction factor 0.23. The mean of technical duplicates is shown in the growth curves.

### Whole-genome sequencing

Genomic DNA was extracted from 2 - 4 mL of overnight culture in LB 0.25 mM DAP, using a kit from Macherey Nagel. A minimum of 300 ng DNA was sent for sequencing to NovoGene. Results were analyzed using the open source breseq software (Barrick et al., 2014).

### Whole-plasmid sequencing

Plasmids were extracted from strains using the the GeneJET Plasmid-Miniprep-Kit from Thermo Fisher Scientific. Whole-plasmid sequencing was performed by Plasmidsaurus using Oxford Nanopore technology.

### Whole proteome analysis

Four biological replicates of each strain were cultivated in tubes containing 4 mL relaxing M9 medium. Pellets equivalent to 1 mL OD_600_ = 2 were harvested in mid-exponential phase and washed twice with ice-cold PBS. Pellets were stored at -70°C until being processed further. After thawing, the cells were washed three times with ice-cold PBS (15,000 g, 10 min, 4°C) and resuspended in 300 μl lysis buffer (2% sodium lauroyl sarcosinate (SLS), 100 mM ammonium bicarbonate). Then samples were heated for 10 min at 90°C. The amount of proteins was determined by bicinchoninic acid protein assay (Thermo Scientific). Proteins were reduced with 5 mM Tris(2-carboxyethyl) phosphine (Thermo Fischer Scientific) at 90°C for 15 min and alkylated using 10 mM iodoacetamid (Sigma Aldrich) at 20°C for 30 min in the dark. For tryptic digestion 50 µg protein was incubated in 0.5% SLS and 1 µg of trypsin (Serva) at 30°C over night.

After digestion, SLS was precipitated by adding a final concentration of 1.5% trifluoroacetic acid (TFA, Thermo Fischer Scientific). Peptides were desalted by using C18 solid phase extraction cartridges (Macherey-Nagel). Cartridges were prepared by adding acetonitrile (ACN), followed by equilibration with 0.1% TFA. Peptides were loaded on equilibrated cartridges, washed with 5% ACN and 0.1% TFA containing buffer and eluted with 50% ACN and 0.1% TFA, and finally dried in a SpeedVac system.

Dried peptides were reconstituted in 0.1% Trifluoroacetic acid and then analyzed using liquid-chromatography-mass spectrometry carried out on an Exploris 480 instrument connected to an Ultimate 3000 RSLC nano and a nanospray flex ion source (all Thermo Scientific). Peptide separation was performed on a reverse phase HPLC column (75 μm x 42 cm) packed in-house with C18 resin (2.4 μm; Dr. Maisch). The following separating gradient was used: 94% solvent A (0.15% formic acid) and 6% solvent B (99.85% acetonitrile, 0.15% formic acid) to 25% solvent B over 40 minutes, and an additional increase of solvent B to 35% for 20min at a flow rate of 300 nl/min. Peptides were ionized at a spray voltage of 2.3 kV, ion transfer tube temperature set at 275 °C, 445.12003 m/z was used as internal calibrant.

MS raw data was acquired on an Exploris 480 (Thermo Scientific) in data independent acquisition (DIA) mode with a method adopted from (Bekker-Jensen et al., 2020). The funnel RF level was set to 40. For DIA experiments full MS resolutions were set to 120.000 at m/z 200 and full MS, AGC (Automatic Gain Control) target was 300% with an IT of 50 ms. Mass range was set to 350–1400. AGC target value for fragment spectra was set at 3000%. 45 windows of 14 Da were used with an overlap of 1 Da. Resolution was set to 15,000 and IT to 22 ms. Stepped HCD collision energy of 25, 27.5, 30 % was used. MS1 data was acquired in profile, MS2 DIA data in centroid mode.

Analysis of DIA data was performed using the DIA-NN version 1.8 (Demichev et al., 2020) using a uniprot protein database from *E*.*coli* including additionally added sequences to generate a data set specific spectral library for the DIA analysis. The neural network based DIA-NN suite performed noise interference correction (mass correction, RT prediction and precursor/fragment co-elution correlation) and peptide precursor signal extraction of the DIA-NN raw data. The following parameters were used: Full tryptic digest was allowed with two missed cleavage sites, and oxidized methionines and carbamidomethylated cysteins. Match between runs and remove likely interferences were enabled. The precursor FDR was set to 1%. The neural network classifier was set to the single-pass mode, and protein inference was based on genes. Quantification strategy was set to any LC (high accuracy). Cross-run normalization was set to RT-dependent. Library generation was set to smart profiling. DIA-NN outputs were further evaluated using the SafeQuant script modified to process DIA-NN outputs (Ahrné et al., 2013; Glatter et al., 2012).

### Lysate activity assay

HOB variant strains were inoculated from cryo stocks into 10 ml LB 0.25 mM DAP, 100 µg/ml streptomycin. After overnight cultivation at 37 °C, they were passaged into 500 ml LB 0.25 mM DAP, 100 µg/ml streptomycin (inoculum 1 ml) and grown at 37 °C overnight. The next day, cells were spun down in a Beckmann Coulter Avanti JXN-26 centrifuge at 4,000 rpm, 10 °C. The resulting pellets were resuspended in 20 ml buffer (50 mM HEPES-KOH pH 7.8, 150 mM KCl) and lysed in an Avestin EmulsiFlex®-B15 High-Pressure Microfluidizer (one passage at 24,000 PSI). Total protein concentration was determined with Roti®Quant reagent (Carl Roth) according to manufacturer’s instructions. Formaldehyde production was observed in reactions containing 100 mM HEPES-KOH pH 7.5, 10 mM MgCl2, 2.5 mM ATP, 250 µM NADPH, 1 mg (lysate only, lysate background) or 0.5 mg lysate (when supplemented with EcAckA or DaArgC) and 100 mM ammonium formate. 0.5 mg DaArgC was added when the EcAckA activity was to be determined instead of full lysate activity. In the same fashion, DaArgC activity was determined by addition of 0.5 mg EcAckA. Reactions were run at 37 °C. Timepoints were taken at 1, 5, 10, 20, 30 and 60 min, where 5 µl samples were quenched in 5 µl 10% formic acid. After all timepoints were taken, 10 µl Nash reagent (2 M ammonium acetate, 20 mM acetylacetone, 50 mM acetic acid) was added and the mixture then incubated at 37 °C for 1 h. For quantification, a 12-step 1:1:1.5 formaldehyde dilution series (starting from 250 µM) was produced and treated the same way. After incubation, the yellow color of the Nash reagent was detected in a TECAN Infinite2000 by its fluorescence at Ex412nm/Em505nm. Initial velocities were determined by linear regression in GraphPad Prism 10.1.0.

### Formaldehyde detection via NASH assay

Formaldehyde in the supernatant of growing cells was detected by reaction with the Nash reagent: 2M ammonium acetate, 20mM acetyl acetone and 50mM acetic acid. In brief, strains were inoculated to an OD of 0.01 in relaxing medium containing 60 mM Na-formate and cultivated in closed tubes at 37°C shaking at 220 rpm. In mid-exponential, 300 uL of culture was harvested and centrifuged at 6500 g for 3 min. The supernatant was transferred to a new tube and used for the NASH assay. For this, samples were mixed 1:1 with the reagent and incubated at 37 °C. After incubation for 1 h, the brightly yellow diacetyldihydrolutidine was formed, which was detected by fluorescence measurements at 412 nm excitation/505 nm emission in 96-well plates in a Spark M10 reader (Tecan; Männedorf, Switzerland). Data was collected using Tecan iControl v3.8.2.0 and Microsoft Excel (v2208). Data analysis and formaldehyde quantification against a formaldehyde calibration curve were performed using GraphPad Prism v9.0.2, where relevant, a linear fit of the individual measurements was applied using the standard settings of the GraphPadPrismtool.

### Data availability

Additional information on the experimental setup as well as detailed results are available from the corresponding author upon request. Proteomics data are available via ProteomeXchange with identifier PXD058946. Any strains and plasmids generated during this study are available upon completing a Materials Transfer Agreement.

