## Supplementary Figures for "Evolution-assisted engineering of formate assimilation via the formyl phosphate route in *Escherichia coli*"

**Supplementary Figures: Evolution-assisted retro-engineering of formate assimilation via the new-to-nature formyl phosphate route in Escherichia coli**

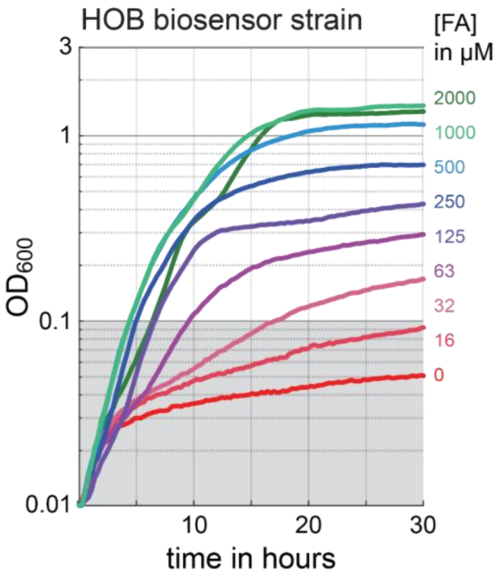


**Figure S1: Formaldehyde tolerance of the HOB biosensor strain.**

The HOB biosensor strain was cultivated in minimal medium supplemented with trace elements, 10 mM glucose, 1 mM isoleucine, 1 mM threonine, 0.25 mM DAP, 50 µM MnCl2, and different concentrations of formaldehyde. Growth was measured over time in a 96 well plate showing a clear dependency of the strain on formaldehyde and a tolerance of up to 2 mM formaldehyde. FA – formaldehyde.

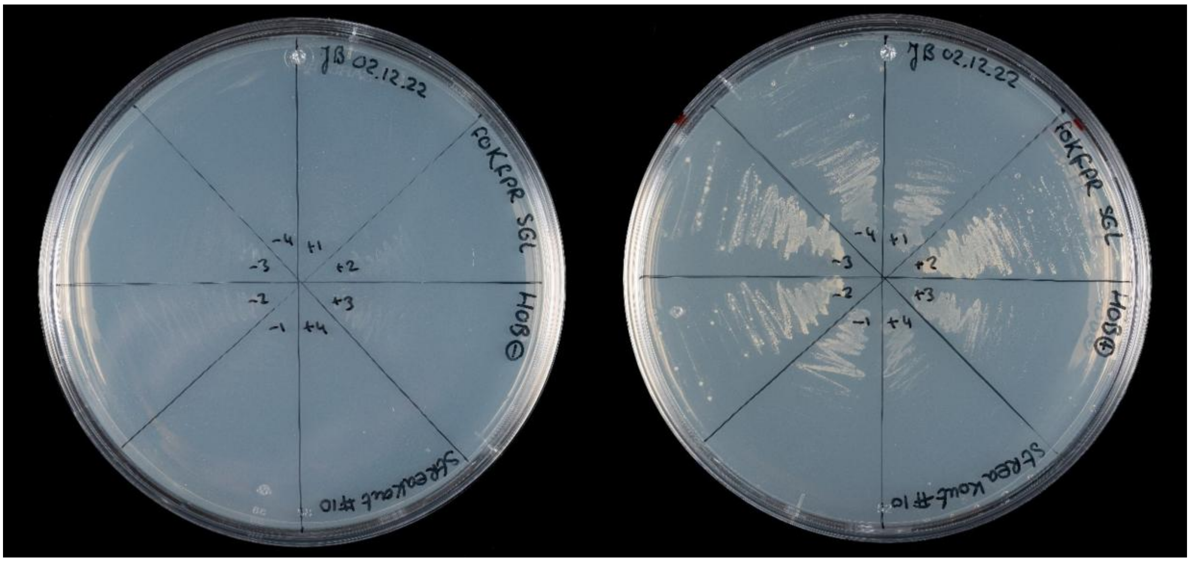


**Figure S2: Plate evolution enabled formate-dependent growth.**

After electroporation with plasmids encoding the formate reduction cascade (ACK-FPR), the HOB biosensor strain was directly plated on minimal medium plates supplemented with 10 mM glucose and 60 mM formate. Small colonies that appeared after two weeks of incubation were streaked over to fresh minimal medium plates supplemented with glucose and formate on a weekly basis to enrich for mutants capable of growing via the formyl phosphate route. The evolved populations were unable to grow on minimal medium plates supplemented with only glucose (left plate) but could grow on minimal medium supplemented with glucose and formate (right plate).

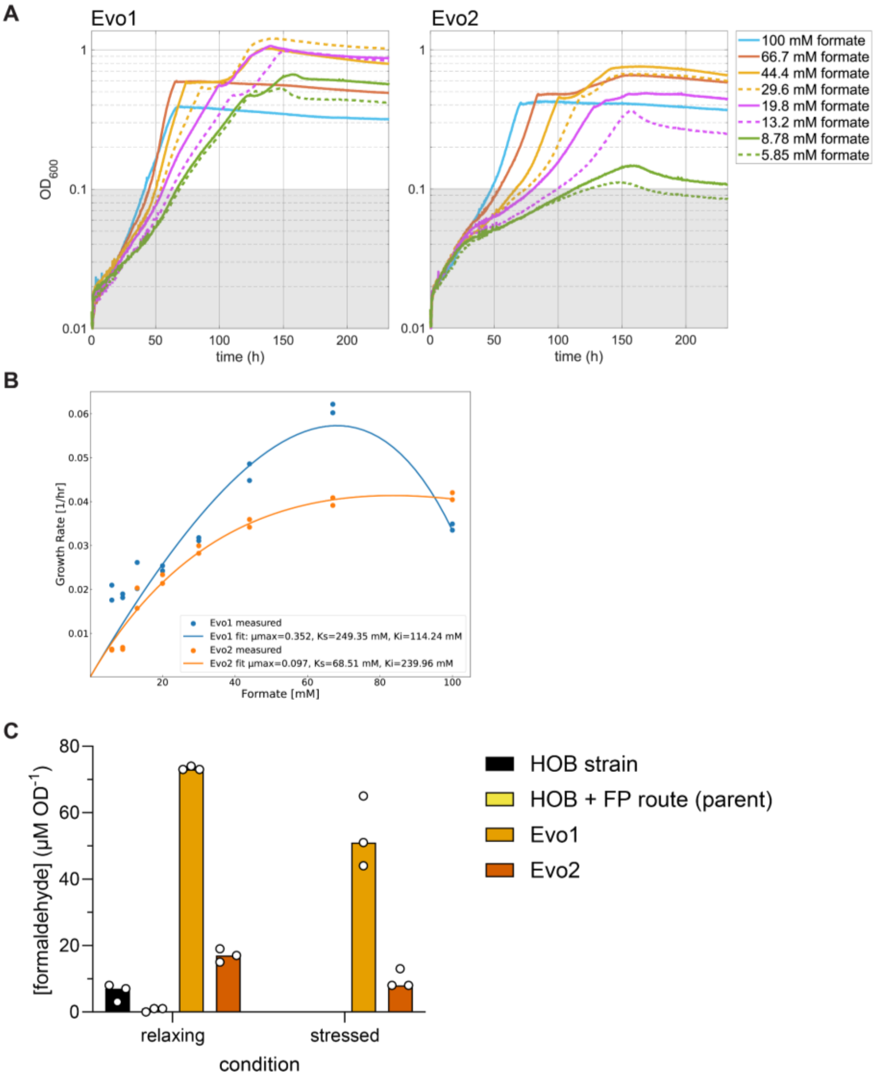


**Figure S3: Analysis of formate-dependent growth of evolved strains.**

(A) The evolved strains Evo1 and Evo2 were cultivated in a defined medium across a gradient of formate concentrations. Both strains exhibited clear formate-dependent growth. (B) From the specific growth rates at each formate level, we estimated kinetic parameters. To characterize strain growth despite the confounding effect of substrate toxicity, the Monod growth model was extended to account for substrate inhibition. Substrate toxicity was modeled as uncompetitive substrate inhibition in the following form (https://doi.org/10.1002/bit.260320404):

$$\mu = \frac{\mu_{max} S \left(1 - \frac{S}{K_i}\right)}{S + K_S \left(1 - \frac{S}{K_i}\right)}$$

where S is the substrate concentration (in this case formate), Ki is the concentration of formate above which the cells cannot grow, and Ks is the Monod constant. By fitting the model to experimental data, we were able to estimate μmax, Ki, and Ks for each strain. The fit resulted in the following parameters: Evo1: μmax=0.352, Ki=114.24 mM, and Ks =249.35 mM, Evo2: μmax=0.097, Ki=239.96 mM, and Ks=68.51 mM. Despite its simple form, the extended Monod growth model reflects the phenotypic differences between the strains. As Evo1 has higher ACK expression, its capacity to metabolize formate, and by extension its maximum growth rate, is higher than Evo2. This however results in a higher production of formaldehyde and ATP usage, which could explain the lower Ki observed. (C) Formaldehyde measurements in culture supernatants were substantially higher for Evo1 than for Evo2 supporting the hypothesis that Evo1 evolved a more active formate reduction cascade.

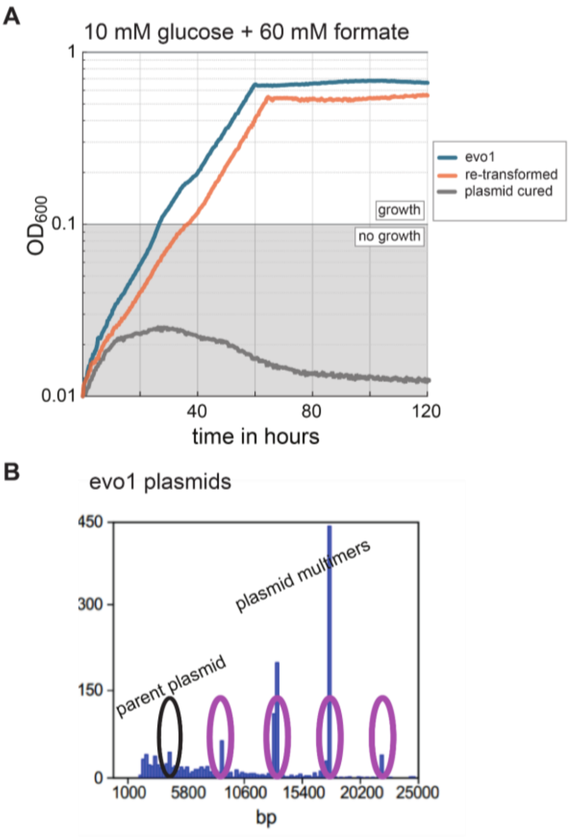


**Figure S4: Plasmid evolution enabled formate-dependent growth.**

(A) The evolved strain Evo1 capable of growing via the formyl phosphate route was cured of its plasmid. Subsequently, the plasmids isolated from Evo1 were transformed into a naive HOB ΔackA-pta strain. Only strains carrying plasmids from the evolved strain were able to grow, indicating that plasmid evolution enabled growth of the evolved strains. (B) Nanopore sequencing of plasmids isolated from Evo1 showed clear signs of plasmid multimerization and a mutation in the regulatory region of ackA pointing towards an adaptation in expression levels of the pathway enzymes.

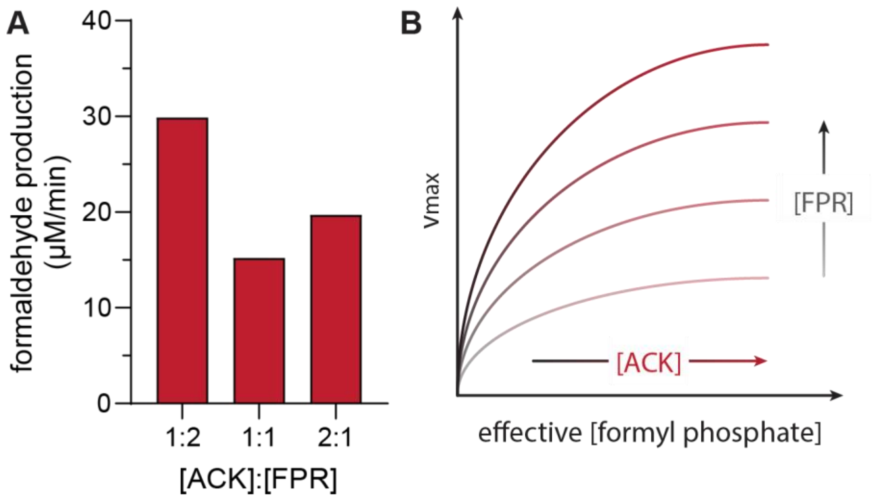


**Figure S5: The influence of [ACK] and [FPR] on cascade productivity.**

(A) In vitro, as previously observed in lysate, the productivity of the cascade is increased both by supplementation of ACK and FPR. Doubling in [FPR] doubles productivity, while doubling [ACK] only yields approximately 33% increase in activity. Assays were performed at 30 °C. Reactions contained 100 mM MOPS pH 7.0, 10 mM MgCl2, 5 mM ATP, and 250 µM NADPH. Enzymes were used at ratios of [ACK]:[FPR] of 2 µM:2 µM, 4 µM:2 µM, and 2 µM:4 µM. Reactions were started by addition of 200 mM NH4 formate. Shown are individual measurements. (B) From these observations, we suggest that FPR is rate-limiting due to its low turnover, impacting cascade productivity in a multiplicative fashion. This is illustrated as a shift of the productivity curves vertically on the Y axis. Increased [ACK], on the other hand, improves cascade performance by increasing formyl phosphate supply. FPR has a high Km on formyl phosphate, meaning it is likely undersaturated in the cascade. Thus, its reaction speed will improve with increasing substrate concentration. This is illustrated by the reaction speed increasing along the X axis, where formyl phosphate supply and [ACK] correlate.

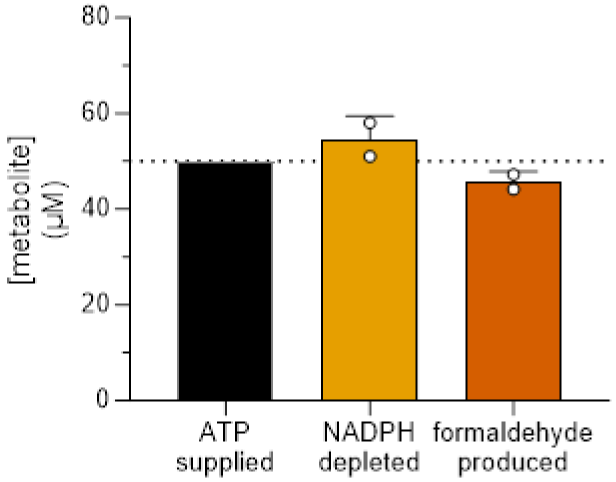


**Figure S6: The ACK-FPR cascade produces stoichiometric amounts of formaldehyde when limited by ATP, and thus is not impacted by formyl phosphate hydrolysis.** Enzymes were supplied with excess NADPH and formate, but limited by 50 µM ATP. Both the depletion of NADPH and formation of formaldehyde are observed to be stoichiometric, suggesting that there is no impact of formyl phosphate hydrolysis on cascade performance. Assays were performed at 30 °C. Reactions contained 100 mM HEPES-KOH pH 7.5, 10 mM MgCl2, 50 µM ATP, 250 µM NADPH, and 5 µM of each enzyme. Reactions were started by addition of 200 mM NH4 formate. Depletion of NADPH was assayed by tracking ABS340nm in a CARY Spectrophotometer. Production of formaldehyde was assessed by NASH assay. Bars represent mean of technical duplicates. Error bars indicate standard deviation. Individual replicates are shown.
