## Supplementary Tables for "Evolution-assisted engineering of formate assimilation via the formyl phosphate route in *Escherichia coli*"

**Supplementary Tables: Evolution-assisted engineering enables Escherichia coli to grow via a new-to-nature formate reduction cascade**

Table S1: List of strains and plasmids used in this study.

| Strain name | Genotype | Plasmid | Source |
|---|---|---|---|
| DH5α | *$F^-$ $\lambda^-$ Φ80lacZΔM15 Δ(lacZYA-argF)U169 deoR recA1 endA1 hsdR17(rK$^-$ mK$^+$) phoA supE44 thi-1 gyrA96 relA1* | | (Hanahan, 1983) |
| MG1655 | *$F^-$ $\lambda^-$ ilvG$^-$ rfb-50 rph-1* | | (Blattner et al., 1997) |
| SIJ488 | MG1655 *Tn7::para-exo-beta-gam; prha-FLP; xylSpm-IsceI* | | (Jensen et al., 2015) |
| HOB biosensor | *ΔfrmRAB ΔpatZ Δasd ΔrhmA*<br>*SS2:Ps:rbsA:rhmA* | | (Schann et al., 2024) |
| HOB Δ*ackA-pta* | *ΔfrmRAB ΔpatZ Δasd ΔrhmA ΔackA-pta*<br>*SS2:Ps:rbsA:rhmA* | | This study |
| HOB Δ*ackA-pta* p:ACK-FPR<br>(Parent) | *ΔfrmRAB ΔpatZ Δasd ΔrhmA ΔackA-pta*<br>*SS2:Ps:rbsA:rhmA* | *p15A:$P_{pgi-20}$: rbsC:ackA:rbsA:NoHis_argC_S178V_G182V_L233I: $Strep^R$* | This study |
| HOB Δ*ackA-pta* p:ACK-FPR evo 1<br>(Evo 1) | *ΔfrmRAB ΔpatZ Δasd ΔrhmA ΔackA-pta*<br>*SS2:Ps:rbsA:rhmA* | *p15A:$P_{pgi-20}$: rbsC*:ackA:rbsA:NoHis_argC_S178V_G182V_L233I:$Strep^R$* | This study |
| HOB Δ*ackA-pta* p:ACK-FPR evo 2<br>(Evo 2) | *ΔfrmRAB ΔpatZ Δasd ΔrhmA ΔackA-pta*<br>*SS2:Ps:rbsA:rhmA* | *p15A:$P_{pgi-20}$: rbsC:ackA*:rbsA:NoHis_argC_S178V_G182V_L233I:$Strep^R$* | This study |
| HOB Δ*ackA-pta* SS9:Ps:A:ackA | *ΔfrmRAB ΔpatZ Δasd ΔrhmA ΔackA-pta*<br>*SS2:Ps:rbsA:rhmA SS9:Ps:rbsA:ackA* | | This study |
| HOB Δ*ackA-pta* SS9:Ps:A:ackA<br>p:FPR1<br>(FR1) | *ΔfrmRAB ΔpatZ Δasd ΔrhmA ΔackA-pta*<br>*SS2:Ps:rbsA:rhmA SS9:Ps:rbsC:ackA* | *p15A:$P_{pgi-20}$: rbsC*:ackA:rbsA:NoHis_argC_S178V_G182V_L233I: $Strep^R$* | This study |
| HOB Δ*ackA-pta* SS9:Ps:A:ackA<br>p:FPR2<br>(FR2) | *ΔfrmRAB ΔpatZ Δasd ΔrhmA ΔackA-pta*<br>*SS2:Ps:rbsA:rhmA SS9:Ps:rbsC:ackA* | *pMB1:$P_{pgi-20}$: rbsC*:ackA:rbsA:NoHis_argC_S178V_G182V_L233I: $Strep^R$* | This study |

Table S2: List of mutations observed in the HOB biosensor and the two evolved strains Evo1 and Evo2 via whole genome sequencing

| **location** | **position** | **mutation** | **HOB biosensor** | **Evo1** | **Evo2** | **gene** | **description** |
|---|---|---|---|---|---|---|---|
| **Chromosome** | 792,844 | A→G | 100% | 100% | 100% | *aph(3')-II (or nptII)* → | aminoglycoside phosphotransferase from Tn5 |
| **Chromosome** | 1,301,174 | Δ1,199 bp | 100% | ? | ? | *insH21* | *insH21* |
| **Chromosome** | 1,621,105 | G→T | 100% | 100% | 100% | *marR* → | DNA-binding transcriptional repressor MarR |
| **Chromosome** | 2,422,165 | A→T | 100% | 100% | 100% | *rpnB* ← | recombination-promotin g nuclease RpnB |
| **Chromosome** | 2,940,367 | G→C | 100% | 100% | 100% | *gcvA* ← | DNA-binding transcriptional dual regulator GcvA |
| **Chromosome** | 4,124,193 | Δ1,200 bp | 100% | ? | 100% | *insH-5* | *insH-5* |
| **Chromosome** | 4,303,486 | +GC | 100% | 100% | 100% | *gltP* → / ← *yjcO* | glutamate/aspartate : H(+) symporter GltP/Sel1 repeat-containing protein YjcO |
| **Plasmid** | 1 | Δ4,411 bp | 100% | | | | plasmid not present in HOB biosensor |
| **Plasmid** | 945 | Δ4 bp | | 100% | 100% | *unknown* ← / → *D aArgC* | Promoter mutation |
| **Plasmid** | 1,011 | A→C | | 100% | | *unknown* ← / → *D aArgC* | RBS C point mutation |
| **Plasmid** | 1,035 | +ATCA | | | 100% | *unknown* ← / → *D aArgC* | 4 bp insertion in AckA: ATGA at position 1036 (17 bp into ackA) leading to the use of an alternative start codon and resulting in an AckA missing the HIS tag |
| **Plasmid** | 2,840 | 2 bp→GT | | 100% | 100% | *DaArgC* → | S178V |
| **Plasmid** | 2,853 | 2 bp→TT | | 100% | 100% | *DaArgC* → | G182V |
| **Plasmid** | 3,005 | C→A | | 100% | 100% | *DaArgC* → | L233I |
